# A complete morphological characterization of all life stages of the phorid fly *M. scalaris*

**DOI:** 10.1101/2023.05.18.541282

**Authors:** Jayakumar Pallavi, Harshita Snehal, Daniela Salazar, Mrunal Hanbar, Larina Bejoy Chiramel, Khushi Alok Jha, Rakshita Sukruth Kolipakala, Sai Bhumica Lakshmi Venkatesh, Tanishka Dayanand Shetty, Navya Madhusudan, Amrutha Mohan, Amulia John, Naomi Deep D’souza, Priyanka Sheet, Deepesh Nagarajan

## Abstract

*Megaselia scalaris*, commonly known as the scuttle fly, is a cosmopolitan species in the family Phoridae. It is an easily cultured fly species that is an emerging model organism in the fields of genetics and developmental biology. Its affinity for carrion and its predictable life cycle makes it useful in the field of forensic science for estimating the post-mortem interval (PMI) of human remains. Cases of human myasis caused by *M. scalaris* have also been reported in the medical literature. Despite its ubiquitous prevalence and its relevance across multiple fields, its morphology has not been adequately characterized. Here, we report the complete morphological characterization of all lifestages of *M. scalaris*, ranging from egg to adult. Scanning electron microscopy has enabled us to uncover morphological features and developmental processes that have previously not been reported in the literature. Our data lays the groundwork for future genetic studies: a morphological characterization of the wild type must be performed before mutants displaying different phenotypes can be identified. In this vein, we also report the observation of a acephalic, or ‘headless’, adult phenotype whose study could yield insights into the process of cephalogenesis.

## Introduction

*Megaselia scalaris*, commonly known as the “scuttle fly”, is a member of the subfamily Metopininae and the newly reinstated tribe Gymnophorini. The Phoridae family contains over 225 genera and over 2,500 species^1^. This species can be distinguished from other phorid flies through its characteristic black stripes on its dorsal abdomen, thick costal vein, and humpbacked appearance^2^. Other distinctive features for this species include a shorter and broader sixth tergite on female adults and a single strong bristle on left side of the epandrium on male adult^3^. The life cycle of *M. scalaris* consists of four stages: eggs, larva, pupa, and adult, with the larval stage consisting of three sequential instars. The life cycle spans 1-3 weeks^4^, and is dependant on temperature and humidity. *M. scalaris* larvae can subsist on a variety of food sources, including carrion^5^, bananas^6^, and even paint^7^. *M. scalaris* breeds prodigiously; more than 350 eggs can be laid by a single female^8^. These traits make *M. scalaris* very easy to rear in the laboratory.

*M. scalaris* has been increasingly used as a model organism in the fields of genetics and developmental biology^8^. Wolf *et al*. used *M. scalaris* for studying the role of histone acetylation in spermiogenesis^9^, observing that histone acetylation occurs before histone replacement with protamines in spermatid nuclei. *M. scalaris* adults display an unusual style of locomotion for phorid flies: they ‘scuttle’ in short bursts followed by periods of rest. Tretyn *et al*. believe that understanding the physiological basis for this behaviour could advance the field of neuromuscular physiology^10^. Tretyn *et al*. also noted differences between the neuromuscular junctional architecture and electrophysiology of *D. melanogaster* and *M. scalaris* larvae. Orgogozo and Schweisguth^11^ noted differences in the peripheral nervous systems (PNS) of *D. melanogaster* and *M. scalaris*. Such differences across phorid and callophoridae flies (blow flies) were used to trace the evolutionary origins of the PNS. Yoo *et al*.^12^, Smith^13^, and Kwak^14^, independently used *M. scalaris* to study apoptosis.

*M. scalaris* has gained importance in the field of forensic entomology. Determining the post-mortem interval (PMI) of a corpse using developmental data for *M* .*scalaris* was found to be more accurate than the PMI calculated using callophoridae flies, especially for corpses found indoors and in winter conditions. Reibe and Madea^15^ demonstrated this using three corpses autopsied in Germany. Greenberg and Wells^16^ described the isolation of *M* .*scalaris* from two corpses in Chicago, and provided temperature-dependant developmental data for *M. scalaris* larvae.

*M. scalaris* is a clinically relevant organism. It has been reported as the etiological agent in several clinical cases of myasis, such as: urinary myasis (Egypt^17^), urinogenital myasis (India^18^, Iran^19^), intestinal myasis (Egypt^20^), nasopharyngeal and wound myasis (Kuwait^21^).

Given its importance in basic research, forensics, and medicine, the existing literature describing the morphological and ultrastructural characterization of *M. scalaris* remains fragmentary and incomplete. Disney^2^ published an influential treatise dealing with the morphology, ecology, development and identification of phorid flies. Due to the large scope of this work, covering several different species in the family Phoridae, and due its reliance on hand-drawn diagrams rather than micrographs, it cannot be used as (and was not designed as) a comprehensive resource dealing with *M. scalaris* morphology.

Other workers have used electron microscopy to characterize different stages of the *M. scalaris* life cycle. However, this characterization is fragmentary as the work was performed by different teams, using different instruments, using different strains of *M. scalaris strains*, and in different countries, making straightforward comparisons between the data difficult. The egg life stage was characterized by Furukawa *et al*.^22^ in Japan, Greenberg and Wells^16^ in Chicago, and Wolf *et. al*^23^ using the “Wienn” strain. The larval life stages were characterized by Boonchu *et al*.^24^ and Sukontason *et al*.^25^ in Thailand, Ismail^26^, Shaheen *et al*.^27^, and Mayzad *et al*.^28^ in Egypt, and Machkour-M’Rabet *et al*.^29^ in Mexico. The pupal lifestage was characterized by Sukontason *et al*.^30^ in Thailand and Braga *et. al*^31^ in Brazil. Furthermore, the individual reports are sometimes contradictory to each other and to our current work. We have addressed these contradictions in the results section.

In this work, we have used light microscopy (LM) and environmental scanning electron microscopy (ESEM) as complementary tools while characterizing *M. scalaris*. Light microsocopy can reveal pigmentation patterns that ESEM cannot detect. ESEM can reveal detailed surface ultrastructures that light microscopy cannot detect, due to both the limited resolution of photons and the optically translucent nature of our samples. Using these tools, we have characterized the egg, larval, pupal, and adult live stages of *M. scalaris* in detail. We believe this work can serve as a morphological ‘atlas’ to others working in the field. Furthermore, we have observed morphological features, developmental processes, and an acephalic, or ‘headless’, adult phenotype that were previously not reported in the literature.

## Results

*M. scalaris* was isolated in Mumbai, India. The specimen’s identity was confirmed via mitochondrial 16s rRNA (supplementary table S1) and cytochrome c oxidase subunit I (COI) (supplementary table S2) sequencing, performed on undifferentiated larvae by Sakhala Enterprises (Bangalore). The morphology of the egg, first instar larval, second instar larval, third instar larval, pupal, and adult (male and female) lifestages is described below.

### The egg lifestage

The eggs of *M. scalaris* are small, ovoid, and yellow-white (Figure 1A,B). The dorsal surface appears rough, possessing spicula, while the ventral surface appears smooth (Figure 1A). ESEM revealed dorsal spicula of three sizes: large (*α*-spicula), medium (*β* -spicula), and small (*γ*-spicula) (Figure 2A-C). *α*-spicula can be described as tooth-shaped, with a broad base curving upwards. *β* -spicula are smaller, spheroid, and more numerous. *γ*-spicula are the smallest observable structures on the dorsal surface. They are irregularly shaped and are the most numerous spicula observed. ESEM revealed a smooth ventral surface composed of hexagonal tiles (Figure 2D). These tiles possess small protuberances on their surface (Figure 2E), that we term as intrahexagonal papillae.

**Figure 1.**
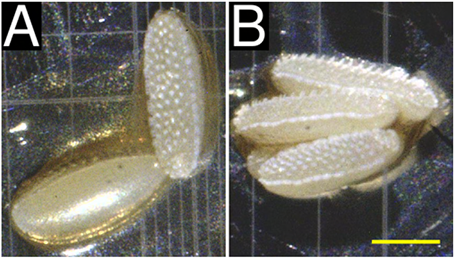
Morphological and morphometric characterization of *M. scalaris* eggs using light microscopy, at 40× magnification. **(A)** Dorsal (above, right) and ventral (below, left) views of eggs. The dorsal surface appears rough due to dorsal spicula that are better visualized using electron microscopy. The ventral surface appears smooth. **(B)** Lateral view of eggs. Dorsal spicula are clearly visualized. A lateral ridge separating the dorsal and ventral surfaces is visible, but is better visualized using electron microscopy. Scale bar (yellow) represents 0.25 mm.

**Figure 2.**
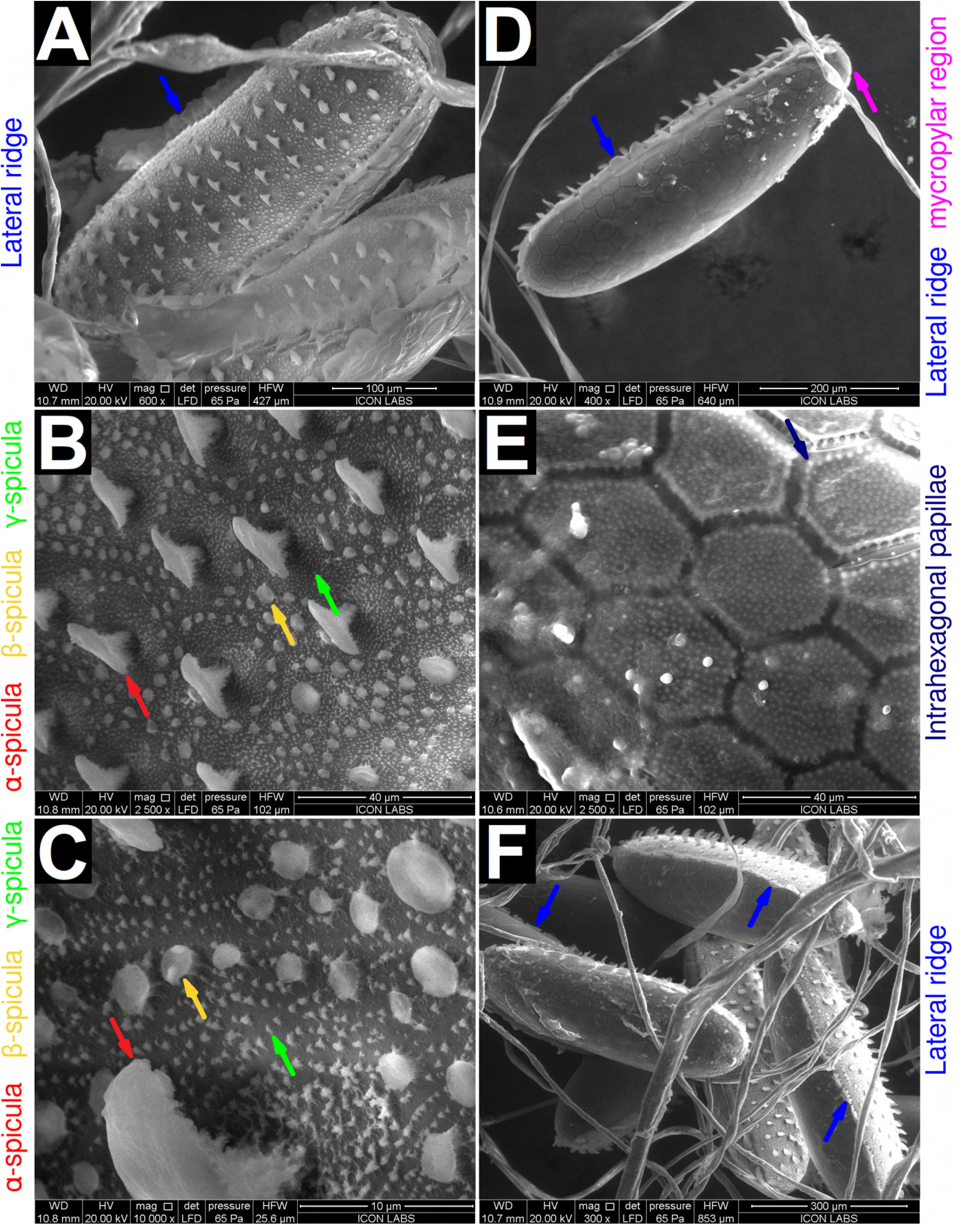
ESEM characterization of *M. scalaris* eggs. **(A)** The dorsal view of a single egg at 600× magnification. Dorsal spicula are visible. Lateral skirt segments (blue arrow) are also visible. **(B)** Dorsal view at 2500× magnification. Three distinct types of spicula are visible: *α*-spicula (red arrow), *β* -spicula (yellow arrow), and *γ*-spicula (green arrow). *α*-spicula are tooth-shaped. *β* -spicula are spheroidal. *γ*-spicula are not well-resolved at this magnification. **(C)** Dorsal view at 10,000× magnification. *γ*-spicula (green arrow) are well resolved at this magnification and appear to be irregular in shape. *α*- and *β* -spicula are also visible (red and yellow arrows, respectively). **(D)** A single egg viewed from the ventral view at 400× magnification. The ventral surface appears to be smooth and covered in a hexagonal tiling. The mycropylar region is visible (magenta arrow). Lateral skirt segments (blue arrow) are also visible. **(E)** Ventral view at 2500× magnification. Papillae within each hexagonal tile are clearly visible. **(F)** Lateral view of multiple eggs at 300× magnification. The lateral skirt (blue arrow) is visible on three separate eggs.

We hypothesize that the dorsal *α*-spicula may help the egg attach to fibrous surfaces via mechanical interlocking. The *β* - and *γ*-spicula may also serve some role in adherence to a substrate. Further, we hypothesize that the papillae on the smooth ventral surface may help attach the egg to hard, smooth surfaces. However, further experiments are required to validate these hypotheses.

A lateral ridge was observed under both light microscopy (Figure 1B) and ESEM (Figure 2A,D,F). This ridge is composed of thin rectangular segments. The mycropylar region is also visible (Figure 2D).

Our observations are mostly in agreement with work published by Furukawa *et al*. on *M. scalaris* specimens isolated in Japan^22^. However, our work offers clearer images acquired at higher resolutions. Similar work was performed by Greenberg and Wells on specimens isolated from a corpse in Chicago, USA^16^. In this case, the eggs displayed extensive structural deformation, possibly occurring during fixation prior to SEM. As we performed ESEM (environmental SEM), our specimens required no treatment prior to visualization.

### The first instar larval lifestage

The larval lifestage of *M. scalaris* can be subdivided into three instars. First instar larvae are characterized by their small size, lack of thoracic or abdominal spiracles, and their lack of pigmentation. No distinct morphological characteristics were observable under light microscopy (Figure 3A,B). However, ESEM revealed a segmented body with three thoracic segments and 8 abdominal segments (Figure 4A), similar to *Drosophila melanogaster* larvae. Delticle bands were observed separating each abdominal segment (Figure 4A,D). Denticle bands are composed of numerous small, electron-reflective (light), toothlike projections. A dorsal view of the larval head (Figure 4B), reveals a bilobed structure possessing a pair of protoantennae in the antennal complex. A ventral view reveals the entire cephalon (Figure 4E) possessing an antennal complex, maxillary palpus, and mouthhooks. A dorsal view of the posterior revealed two immature posterior spiracles (Figure 4C). These structures mature by the second instar, but remain unpigmented. By the third instar, the posterior spiracles develop pigmentation, making them easily observable under light microscopy (Figure 3E). A ventral view of the posterior revealed the anal opening surrounded by anal papillae (Figure 4F). No anal denticula were observed.

**Figure 3.**
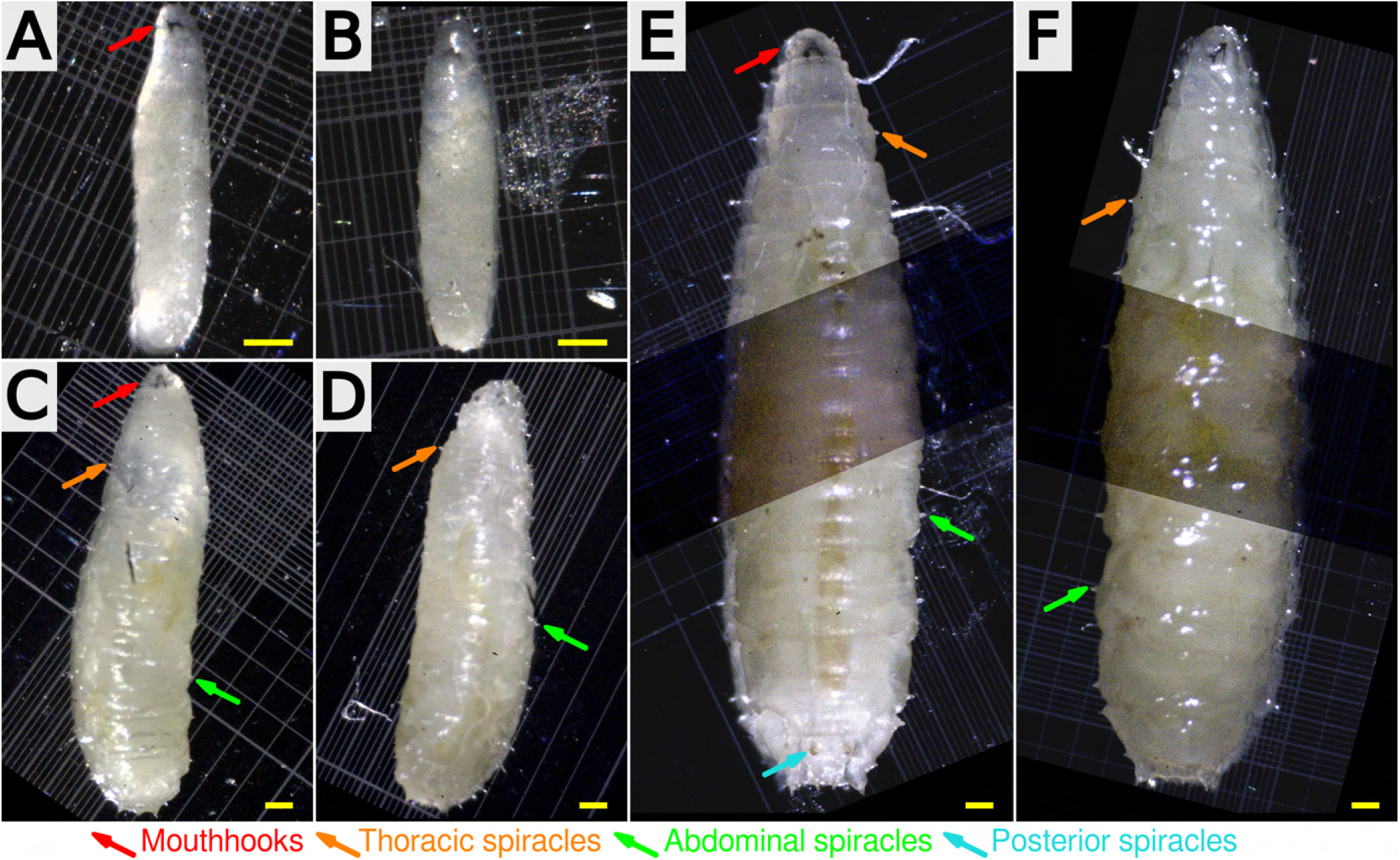
Light microscopic characterization of *M. scalaris* larvae. **First instar larva** at 40× magnification: **(A)** dorsal and **(V)** ventral views. Note the absence of thoracic or abdominal spiracles. Posterior spiracles are immature at this developmental stage (Figure 4C) and cannot be resolved under light microscopy. **Second instar larva** at 20× magnification: **(C)** dorsal and **(D)** ventral views. Posterior spiracles are well-developed at this lifestage (Figure 5E) but are not pigmented. Their lack of contrast makes them difficult to observe under light microscopy. **Third instar larva** at 20× magnification: **(E)** dorsal and **(F)** ventral views. Posterior spiracles are well developed and pigmented (cyan arrow). Thoracic (orange arrows) and abdominal (green arrows) spiracles can be seen under light microscopy for both second instar and third instar larvae. Mouthhooks can be seen in dorsal views of all instars (red arrows). Panels containing third instar larvae are composite images, as the entire specimen could not fit under a single field of view. The dark bands at the centre of the images indicate regions of overlap. Background regions not included in the field of view are colored black. For all panels, the scale bar (yellow) represents 0.25 mm.

**Figure 4.**
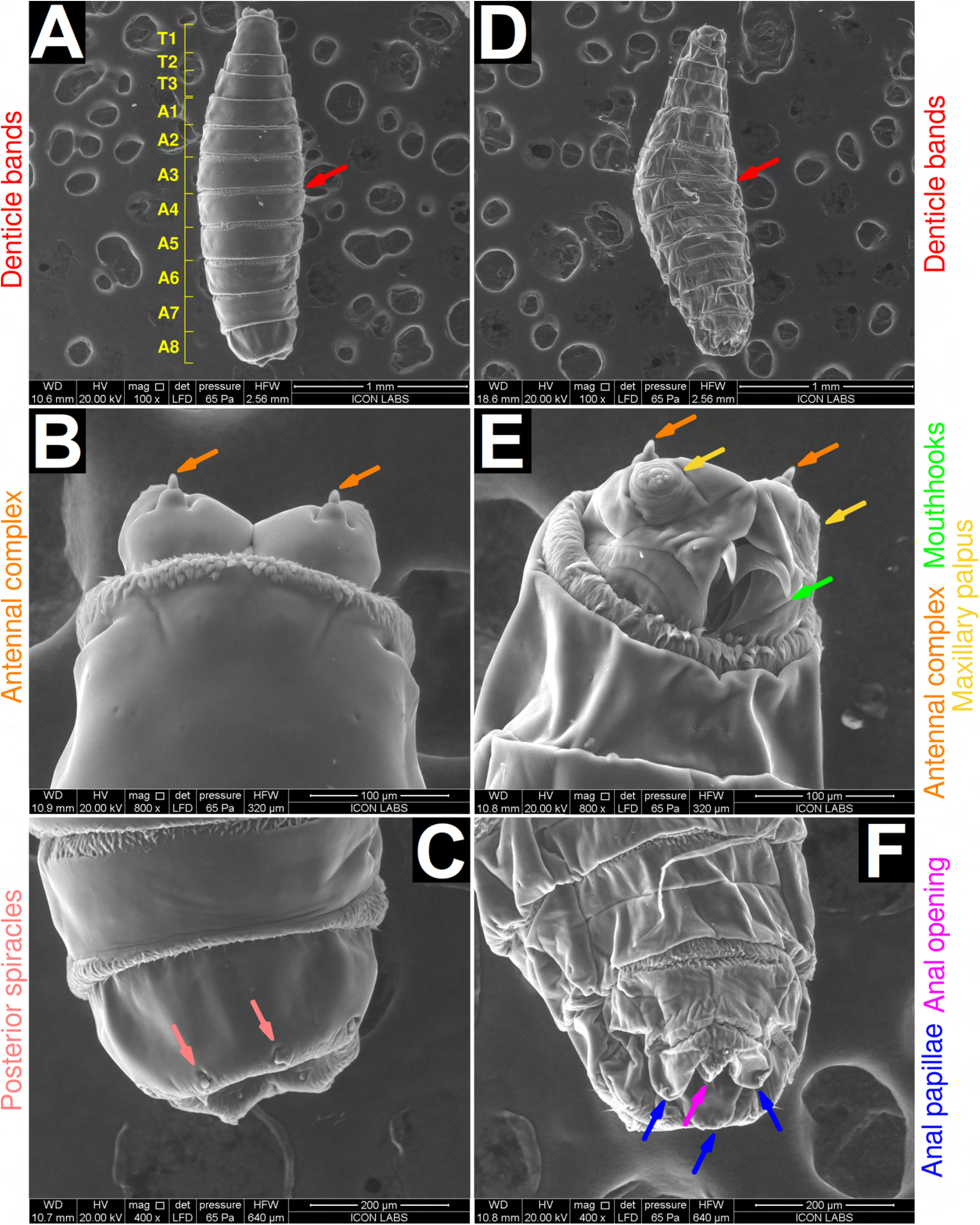
ESEM characterization of *M. scalaris* first instar larvae. **(A)** Dorsal view of a single larvae at 100× magnification. The larva is divided into 12 segments (yellow): three thoracic (T1-T3) and 8 abdominal (A1-A8) segments. Denticle bands (red arrow) are visible in between the abdominal segments. **(B)** Dorsal view of the anterior cephalon. A bilobed stricture possessing two protoantennae (orange arrows) comprising the antennal complex can be seen. **(C)** Dorsal view of the posterior region. Two immature spiracles (pink arrows) can be seen. **(D)** Ventral view of a single larvae at 100× magnification. Isopropanol preservation appears to have denatured this specimen to a greater extent than in the specimen in panels **(A-C)**, nevertheless most morphological features are well preserved. Denticle bands (red arrow) in between the abdominal segments are also seen on the ventral surface. **(E)** Ventral view of the anterior cephalon. The antennal complex (orange arrows), maxillary palpus (yellow arrows) and mouthhooks (yellow arrows) can be seen. The labrum, labium and serrated mouthhooks were not observed. **(F)** Ventral view of the posterior anus. The anal opening (magenta arrow) is surrounded by anal papillae (blue arrows). No anal denticula were observed.

Our observations are in agreement with those of Boonchu *et al*. on first instar larvae isolated in Thailand^24^, except in the following aspects: Boonchu *et al*. did not report the presence of maxillary palpus on the cephalon of first instar larvae. These structures were possibly destroyed during the extensive glutaraldehyde fixation process required to prepare larvae for SEM. As we performed ESEM (environmental SEM), our specimens required minimal treatment and were able to retain fine structures. Boonchu *et al*. reported the presence of labrum and labium on their first instar larval specimen. We did not observe these structures on first instar larvae, but observed them on second and third instar larvae. Boonchu *et al*. reported the presence of semi-circular, lobal, and deeply serrated mouthhooks in their first instar larval specimen. We observed conventional mouthhooks for first instar larvae (Figure 4E). We observed serrated mouthhooks on second and third instar larvae (Figure 5D, 6C,7). Liu and Greenberg^32^ reported conventional mouthhooks on the cephalopharyngeal skeletons of first instar *M. scalaris* larvae, in agreement with our observations.

**Figure 5.**
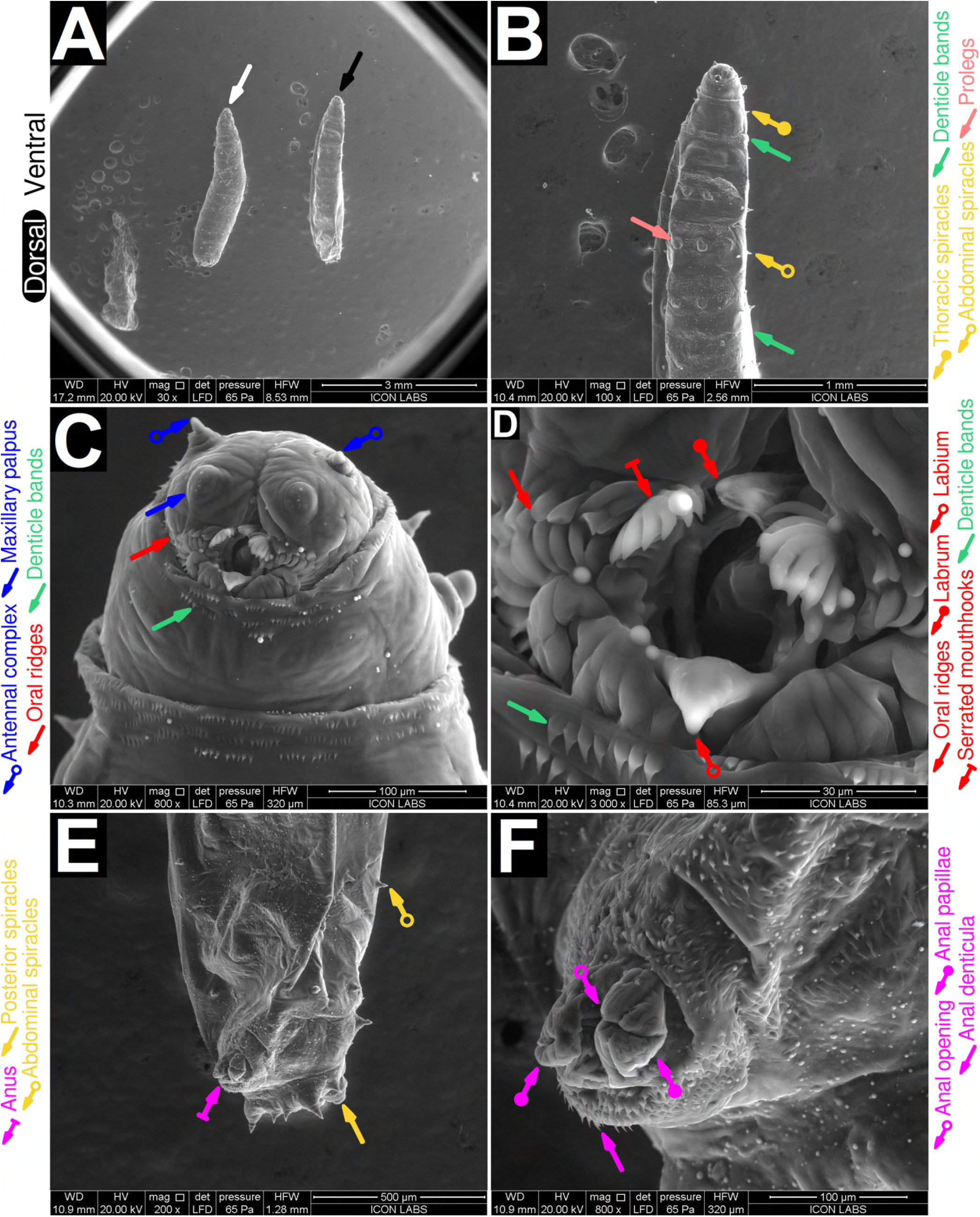
ESEM characterization of *M. scalaris* second instar larvae. **(A)** dorsal (white arrow) and ventral (black arrow) views at 30× magnification. **(B)** Anterior/ventral view of the larva at 100× magnification. The cephalon is seen at the anterior end. Denticle bands (green arrow) are seen in between both thoracic and abdominal segments. Thoracic spiracles (yellow arrow, round end) and abdominal spiracles (yellow arrow, hollow end) can be seen. Prolegs (pink arrow) are present on abdominal segments. **(C)** Anterior view of the cephalon at 800× magnification. Denticle bands (green arrow), oral ridges (red arrow), the maxillary palpus (blue arrow) and the antennal complex (blue arrow, hollow end) can be seen. The rightmost antenna appears embedded, however we believe it to be an artifact of isopropanol preservation **(D)** View of the oral cavity at 3000× magnification. The denticle bands (red arrow), labrum (red arrow, round end), labium (red arrow, hollow end), and serrated moithhooks (red arrow, flat end) can be seen. **(E)** Ventral /posterior view of a larva at 200× magnification. The abdominal spiracles (yellow arrow, hollow end), posterior spiracles (yellow arrow), and anus (magenta arrow, flat end) can be seen. **(F)** View of the anus at 200× magnification. The anal opening (magenta arrow, hollow end), anal papillae (magenta arrow, round end), and anal denticula (magenta arrow) are seen.

### The second instar larval lifestage

Second instar larvae follow the same segmentation pattern as first instar larvae (Figure 5A). Second instar larvae possess a pair of fully developed posterior spiracles (Figure 5E), unlike the immature posterior spiracles observed on first instar larvae (Figure 4C). The posterior spiracles remain unpigmented during the second instar, only developing pigmentation by the third instar (Figure 3E). We observed denticle bands in between both thoracic and abdominal segments (Figure 5B), as well as clusters of denticles all across the body (Fig 5C,D,F). We observed thoracic and abdominal spiracles on every segment of these larvae (Figure 5B,E). Thoracic and abdominal spiracles distinguish second instar larvae from both first instar larvae and larvae of *Drosophila melanogaster*, both of which only possess posterior spiracles. We observed prolegs on abdominal segments (Figure 5B).

We observed a cephalon possessing an antennal complex, maxillary palpus, and a mouth (Figure 5C). The mouth appeared more developed that that of first instar larvae. The mouth possessed oral ridges, a labrum, a labium, and serrated mouthhooks. A large oral cavity can also be seen. The posterior end possessed posterior spiracles and an anus (Figure 5E). The anus itself was comprised of an anal opening and anal papillae (Figure 5F), and was surrounded by a ring of anal denticula, with the individual denticles being similar in morphology to the aforementioned denticle bands.

Boonchu *et al*.^24^ presented electron micrographs of the anterior and posterior ends of second instar larvae, which are in agreement with our observations. However, they did not present thoracic and abdominal electron micrographs. Consequently, discerning any distinctions in the thoracic and abdominal morphology of our specimens in comparison to theirs is unfeasible.

### The third instar larval lifestage

Third instar larvae follow the same segmentation pattern as first and second instar larvae. Third instar larvae possess a pair of fully developed and pigmented posterior spiracles (Figure 3E), unlike the immature posterior spiracles observed on first instar larvae (Figure 4E), or the mature yet unpigmented posterior spiracles found on second instar larvae (Figure 5E). Third instar larvae retain intersegmental denticle bands (Figure 5A,C,D). Thoracic and abdominal spiracles were present on every segment (Figure 6A). Prolegs were present on abdominal segments (Figure 6A).

**Figure 6.**
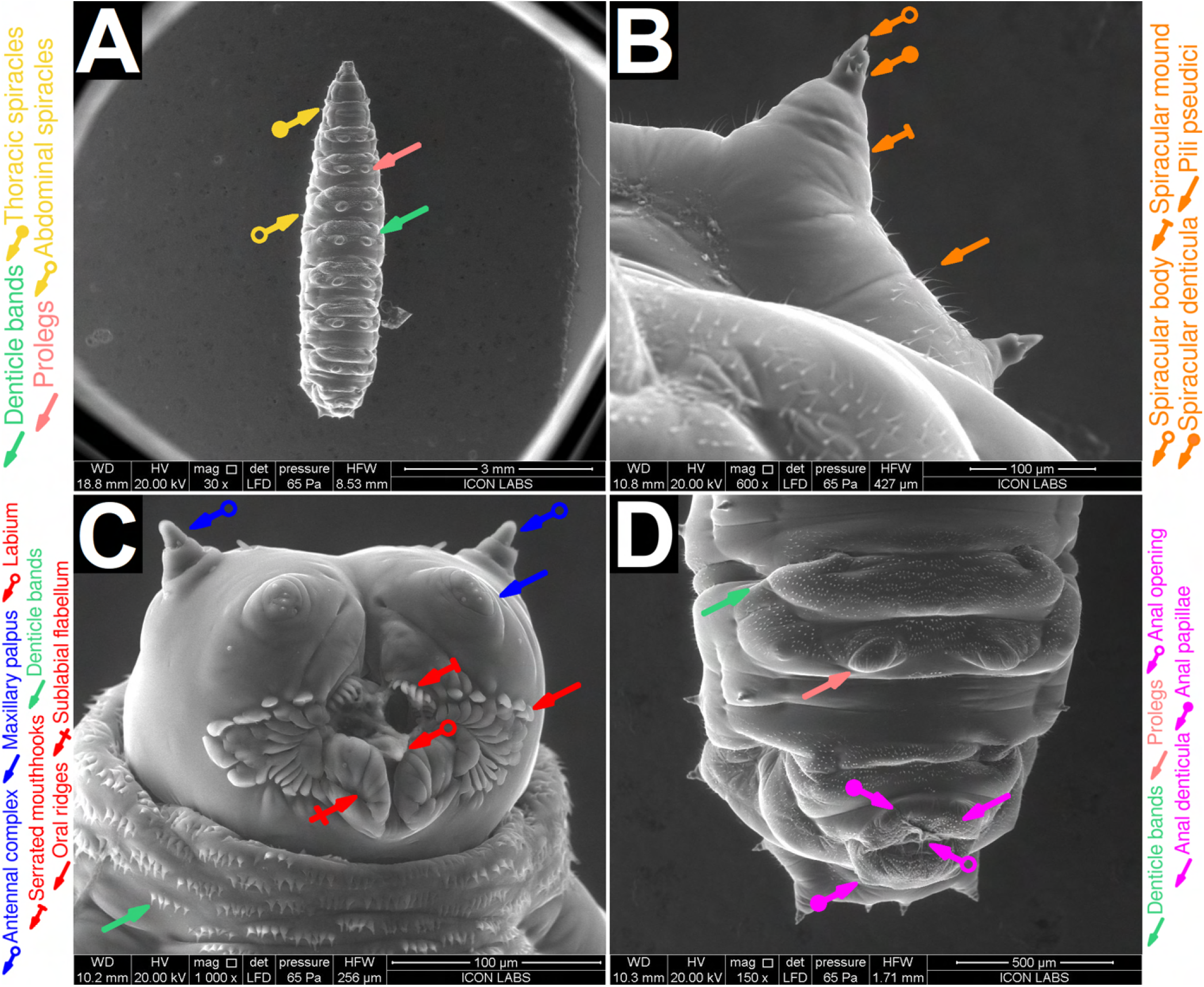
ESEM characterization of *M. scalaris* third instar larvae. **(A)** ventral view at 30*×* magnification. Prolegs (green arrow) are present on every abdominal segment). Thoracic spiracles (yellow arrow, round end) and abdominal spiracles (yellow arrow, hollow end) are present on every segment. Denticle bands (green arrow) are present in between every segment. **(B)** The spiracular complex at 600*×* magnification. This complex consists of a spiracular mound (orange arrow, flat end), spiracular body (orange arrow, hollow end,) and spiracular denticula (orange arrow, round end). Pili pseudici can also be seen. **(C)** Anterior view of the cephalon at 1000*×* magnification. Denticle bands (green arrow), oral ridges (red arrow), the maxillary palpus (blue arrow) and the antennal complex (blue arrow, hollow end) can be seen. The oral cavity constists of the labium (red arrow, hollow end), serrated mouthhooks (red arrow, flat end), and sublabial flabellum (red arrow, crossed end). **(E)** Ventral posterior view at 150*×* magnification. Prolegs on the abdominal segments (pink arrow), the anal opening (magenta arrow, hollow end), anal papillae (magenta arrow, round end), and anal denticula (magenta arrow) are seen.

Mature spiracles are present on third instar larvae (Figure 6B). These spiracles consist of a wide spiracular mound derived from surrounding tissue that supports spiracular body. The spiracular body possesses spiracular denticula similar in morphology to denticula found in denticle bands. The larval body possesses a sparse covering of thin, hair-like structures that we term as pili pseudici.

The cephalon of this specimen (Figure 6C) possessed an antennal complex, maxillary palpus, oral ridges, the labium, serrated mouthhooks, and a pair of structures we term the sublabial flabella. The labrum was either absent or recessed into the oral cavity. In (Figure 6C), the oral cavity appears open, exposing the mouthhook channels. In Figure 7, all specimens appear to have closed oral cavities. We believe the serrated mouthhooks, the labium, and sublabial flabella act analogously to human lips, opening and closing the oral cavity whenever necessary.

**Figure 7.**
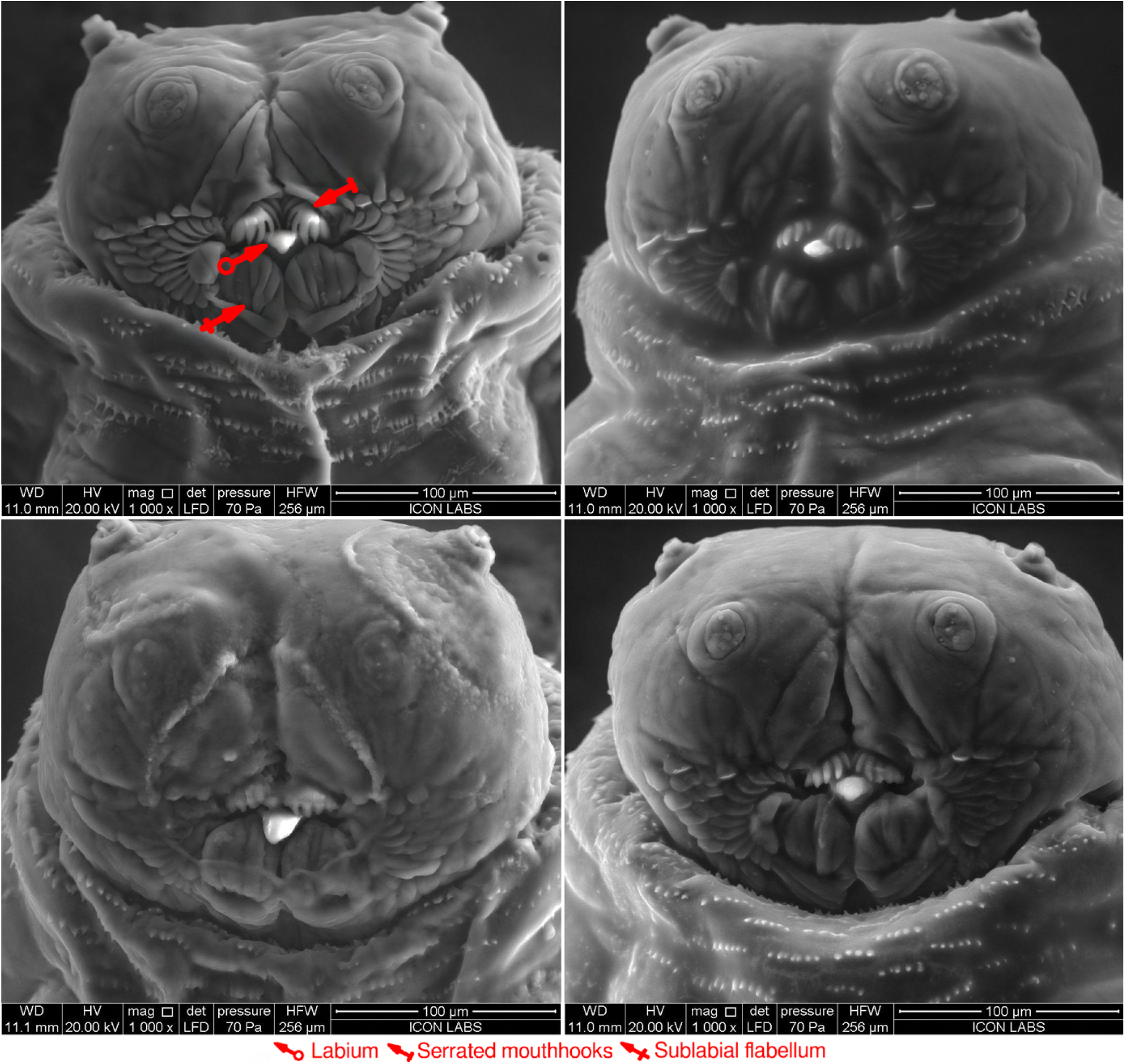
ESEM characterization of the cephalons of four specimens of *M. scalaris* third instar larvae. The labium (red arrow, hollow end), serrated mouthhooks (red arrow, flat end), and sublabial flabellum (red arrow, crossed end) are highlighted for one specimen.

Every third instar larval cephalon we observed possessed a unique pattern of oral ridges (Figure 6C, Figure 7), akin to fingerprints. Our future research will aim to investigate whether the distinct patterns of oral ridges in third instar larvae can be employed as a tool for phylogenetic analysis. We hypothesize that closely related larvae, such as those belonging to the same *M. scalaris* population, will exhibit similar oral ridge patterns, while those from more distant populations will display differing patterns. The anus appeared similar to that of second instar larvae. It possessed an anal opening, anal papillae, and anal denticula (Figure 6D).

Our observations are in agreement with those of Sukontason *et al*.^25^, Ismail^26^, Shaheen *et al*.^27^, and Machkour-M’Rabet *et al*.^29^ on third instar larvae isolated in Thailand, Egypt, Egypt, and Mexico, respectively. However, we have characterized the larval cephalon and spiracular complex in greater detail.

### The pupal life stage

The pupal lifestage of *M. scalaris* can be divided into two distinct sub-stages: the early pupal stage, before the emergence of respiratory horns and the development of the carapace. The later pupal lifestage occurs after these events.

The early pupal stage is characterized by a cylindrical, segmented puparium following the same thoracic and abdominal segmentation pattern as third instar larvae (Figure 8A, Figure 9A). We term the three thoracic segments and three abdominal segments present in the dorso-anterior region as the precarapace (Figure 9A, Figure 8A). A thin septum bisecting the precarapace can be observed. We term this structure as the carapogenic septum (Figure 9A, Figure 8A). The anterior end of the precarapace contains the dorsal cap. The pupa retains its larval spiracular network, which includes the thoracic/abdominal (Figure 9A), anterior 9B), and posterior spiracles 9C). The posterior spiracles possess two straight slits. Denticle bands are retained from the larval lifestage.

**Figure 8.**
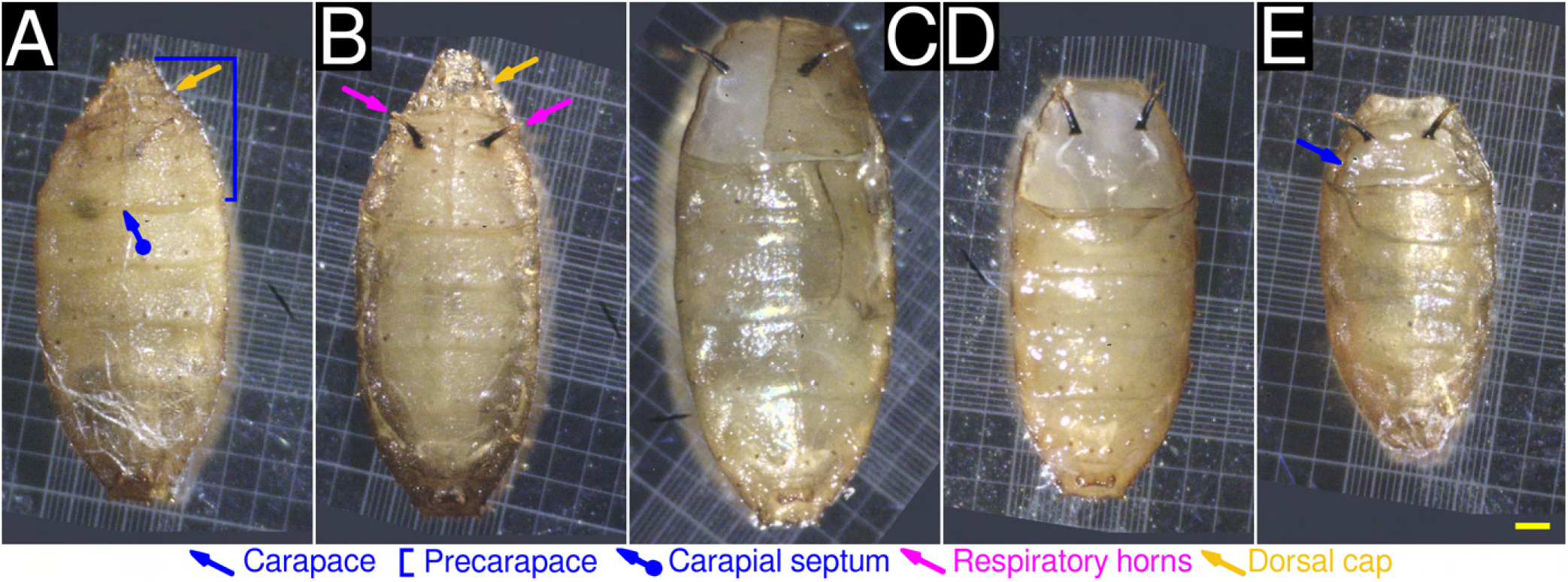
Light microscopic characterization of *M. scalaris* pupae at 20× magnification. **(A)** Dorsal view of a pupa (early stage) displaying the dorsal cap (yellow arrow), precarapace (blue bracket) and carapogenic septum (blue arrow, round end). The respiratory horns are absent in this stage. **(B)** Development of respiratory horns (magenta arrows). **(C)** Ecdysis of one lateral segment of the precarapace along the carapogenic septum. Also note the ecdysis of the dorsal cap. **(E)** Ecdysis of both lateral segments of the precarapace along the carapogenic septum. **(E)** Pupa (late stage) with fully formed carapace (blue arrow). It should be noted that a different specimen is present on each panel. Background regions not included in the field of view are colored dark gray. For all panels, the scale bar (yellow) represents 0.25 mm.

**Figure 9.**
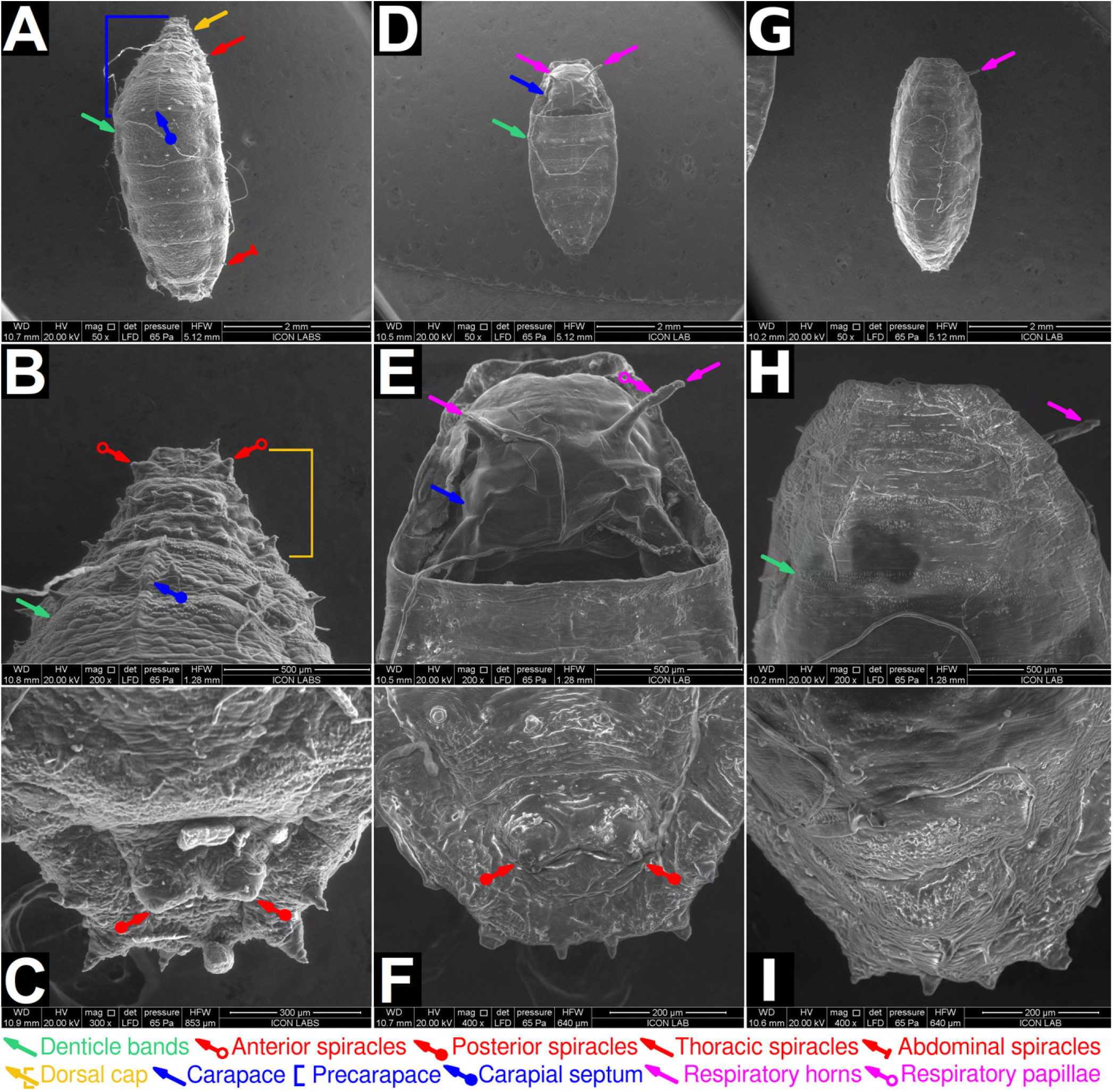
ESEM characterization of *M. scalaris* pupae. **(A)** Dorsal view of a pupa (early stage) at 50× magnification, displaying the dorsal cap (yellow arrow), precarapace (blue bracket), dentice bands (green arrows), thoracic spiracles (red arrow), and abdominal spiracles (red arrow, flat end). The respiratory horns are absent in this stage. **(B)** Pupa (early stage): anterior view (dorsal) at 200× magnification, displaying the dorsal cap (yellow bracket) anterior spiracles (red arrows, hollow end). Spiracluar openings (slits) can be seen. **(C)** Pupa (early stage): posterior view (dorsal) at 300× magnification. Posterior spiracles (red arrows, round end) can be seen, each possessing two spiracular openings (slits). **(D)** Dorsal view of a pupa (late stage) displaying prominent respiratory horns (magenta arrows). The carapace (blue arrow), formed by the fusion of thoracic and abdominal segments, can be seen. **(E)** Pupa (late stage): Anterior view (dorsal) at 200× magnification. The carapace (blue arrow) is in clear view. Two respiratory horns (magenta arrows) are seen, however it should be noted that the left respiratory horn was damaged. Spirally arranged respiratory papillae (magenta arrow, hollow end) are seen. **(F)** Pupa (late stage): posterior view (dorsal) at 400× magnification. The posterior spiracles appear to have atrophied. Spiracular openings (slits) were not observed. **(G)** Ventral view of pupa (late stage) at 50× magnification. **(H)** Pupa (late stage): anterior view (ventral) at 50× magnification. **(I)** Pupa (late stage): posterior view (ventral) at 50× magnification

Pupae in the early stage transition to the late stage through a process we term carapogenesis. Firstly, two respiratory horns emerge on the second abdominal segment of early pupae (Figure 8B). Secondly, the two lateral segments of the precarapace undergo sequential ecdysis along the carapogenic septum, exposing the underlying soft tissue (Figure 8C,D). The respiratory horns remain unaffected. The loss of the dorsal cap occurs simultaneously. Thirdly, a chitinous carapace is synthesized and encapsulates the previously exposed soft tissue (Figure 9D,E, Figure 8E).

The later pupal lifestage occurs after the formation of the respiratory horns and carapace. Spirally arranged respiratory papillae can be seen on the surface of the respiratory horns (Figure 9E). The formation of respiratory horns may be required to compensate for the atrophy of all other spiracles (Figure 9F). The pupa now displays dorso-ventral flattening. The ventral view of the later pupal lifestage is unremarkable (Figure 9D,E,F). These pupal lifestages have not been previously reported in the literature.

An intermediate pupal lifestage of *M. scalaris* has also been characterized using SEM by Sukontason *et. al*^30^, and is mostly in agreement with our observations. Sukontason *et. al* reported pupae possessing respiratory horns but no carapace, similar to the pupa shown in Figure 8B. Abdel-Gawad^33^ characterized late stage *M. scalaris* pupae, and is in agreement with our observations. Braga *et. al*^31^ characterized late stage *M. scalaris* pupae in mummified human remains in Brazil. The pupal specimen presented possessed a carapace but did not possess respiratory horns. Respiratory horns are fragile and may not preserve easily. It should be noted that no other group identified distinct sub-stages in pupal development.

### The adult lifestage

*M. scalaris* adults (Figure 10) exhibit sexual dimorphism. The female (Figure 14A) is larger than the male Figure 11A). The sexes can also be distinguished by their mouthparts and genitalia. Both sexes are morphologically similar in all other respects. To avoid repetition, only the male or female specimen will be described while discussing common morphological characteristics. The *M. scalaris* adult body can be divided into the head, thorax, and abdomen.

**Figure 10.**
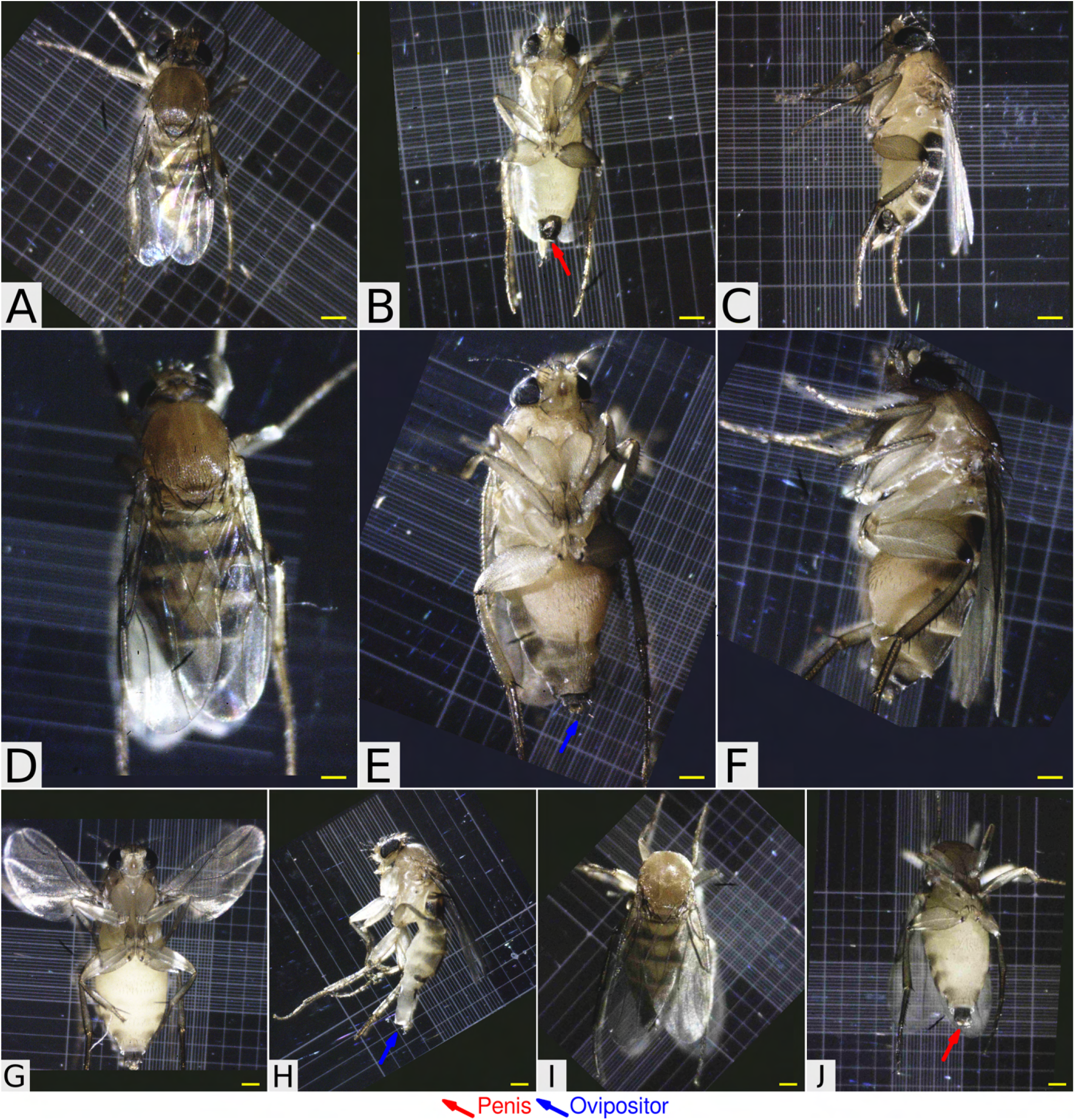
*M. scalaris* adult male and females observed under light microscopy at 20× magnification. For the sake of brevity, descriptions of anatomical features described using ESEM micrographs will not be repeated here. **(A)** Male, dorsal view. Note the characteristic black stripes on every segment of the dorsal abdomen. **(B)** Male, ventral view. The intensely pigmented penis is visible at the posterior end (red arrow). **(C)** Male, lateral view. **(D)** Female, dorsal view. Note the characteristic black stripes on every segment of the dorsal abdomen. **(E)** Female, ventral view. The ovipositor is visible at the posterior end (blue arrow). Female, lateral view. **(G)** Pregnant female. Note the distended abdomen. **(H)** Female immediately after ovipositon. Note the distended ovipositor and reduction in abdominal size. **(I)** Dorsal view of a male displaying the ‘headless’ acephalic phenotype. Ventral view of a male displaying the acephalic phenotype. The male can be identified by the penis (red arrow). Background regions not included in the field of view are colored dark blue. For all panels, the scale bar (yellow) represents 0.25 mm.

**Figure 11.**
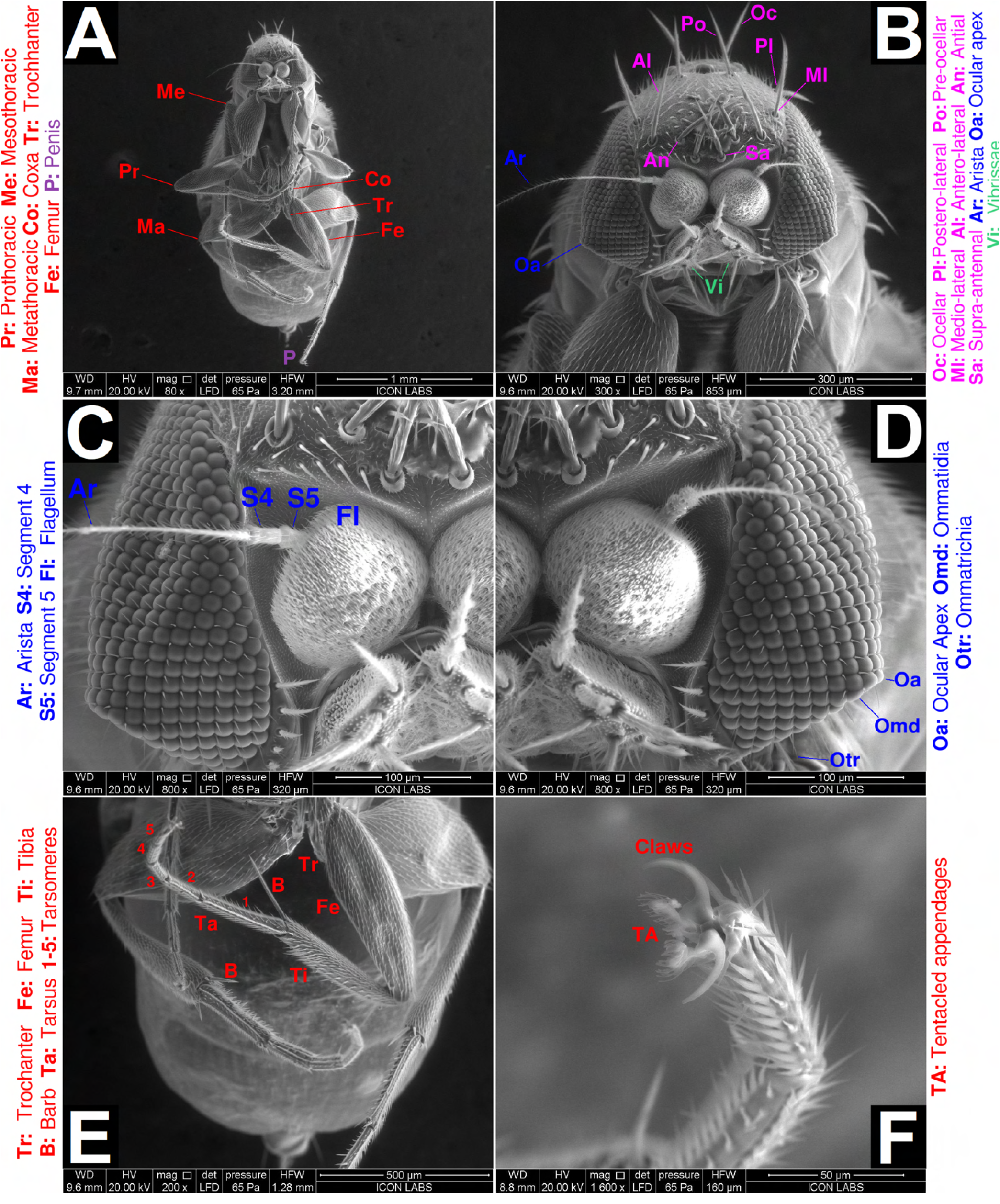
ESEM characterization of adult male *M. scalaris*. **(A)** Ventral view of the body at 80× magnification, displaying a head, thorax, and abdomen (not labelled). Prothoracic (Pr), mesothoracic (Me), metathoracic(Ma) legs are highlighted. The coxa (Co), trochanter (Tr) and femur (Fe) segments of a single mesothoracic leg are highlighted. The penis (P) is present at the posterior end. **(B)** The anterior cephalon at 300× magnification. Ocular (Oc), pseudo-lateral (Pl), pre-ocular (Po), medio-lateral (Ml), antero-lateral (Al), antial (An), and supra-antennal (Sa) bristles are highlighted. The Ocular apex (Oa), arista of the antenna (Ar) and mouthparts termed vibrissae (Vi) are also highlighted. (C,D) Left and right views of the cephalon respectively, at 800× magnification, focusing on the eyes and antennae. The antennal complex consists of the distal arista (Ar), followed by segments 4 and 5 (S4,S5) terminating at the flagellum (Fl). The eyes are composed of numerous ommatidia (Omd) with ommatrichia (otr) present at every interstitial space. The ocular apex (Oa) is also highlighted. Note the hexagonal packing of the ommatidia. (E) Ventral view of the abdomen at 200× magnification. The trochanter (Tr), femue (Fe), tibia (Ti) Tarsus (Ta), and tarsomeres (1-5) for a single mesothoracic leg are highlighted. Barbs (B) projecting out of the tibia for this leg are also highlighted. **(F)** The distal end of a single mesothoracic leg at 1600× magnification, showing the claws and tentacled appendages (TA).

The defining feature of the head are the large compound eyes characteristic of all phorid flies. Each compound eye contains hemispherical ommatidia arranged in hexagonal or square close packing. Ommatidia closer to the periphery display hexagonal close packing (Figure 11C,D,14B), while ommatidia closer to the center display square close packing (Figure 12B). Ommatrichia are present in every interstitial space between ommatidia (Figure 12B). Double ommatrichia are occasionally present (Figure 11D). Both our male and female specimens’ eyes deviated from the conventional spherical shape. Their eyes possessed a ventral, pointed structure we term as the occular apex (Figure 11C,D,14B). It should be noted that the ocular apex has not been reported for *D. melanogaster*^34,35^ or any other scuttle fly species^2^. Interestingly, the ocular apex has not been reported for other strains of *M. scalaris* as well. Sukontason *et. al* reported the ultrastructure of eyes from *M. scalaris* specimes isolated in Thailand^36^. Their specimens did not possess ocular apexes. The possibility that the ocular apex is an electron microscopic artifact can be eliminated since both eyes on our male specimen, as well as both eyes on our female specimen, possess ocular apexes. The possibility that such a structure independently arose 4 times by chance is very remote. This leads us to conclude that *M. scalaris* strains possessing an ocular apex may be geographically restricted within a range that may span anywhere from the Mumbai metropolitan area to the Indian subcontinent. Further work is needed to confirm the range of this phenotype.

**Figure 12.**
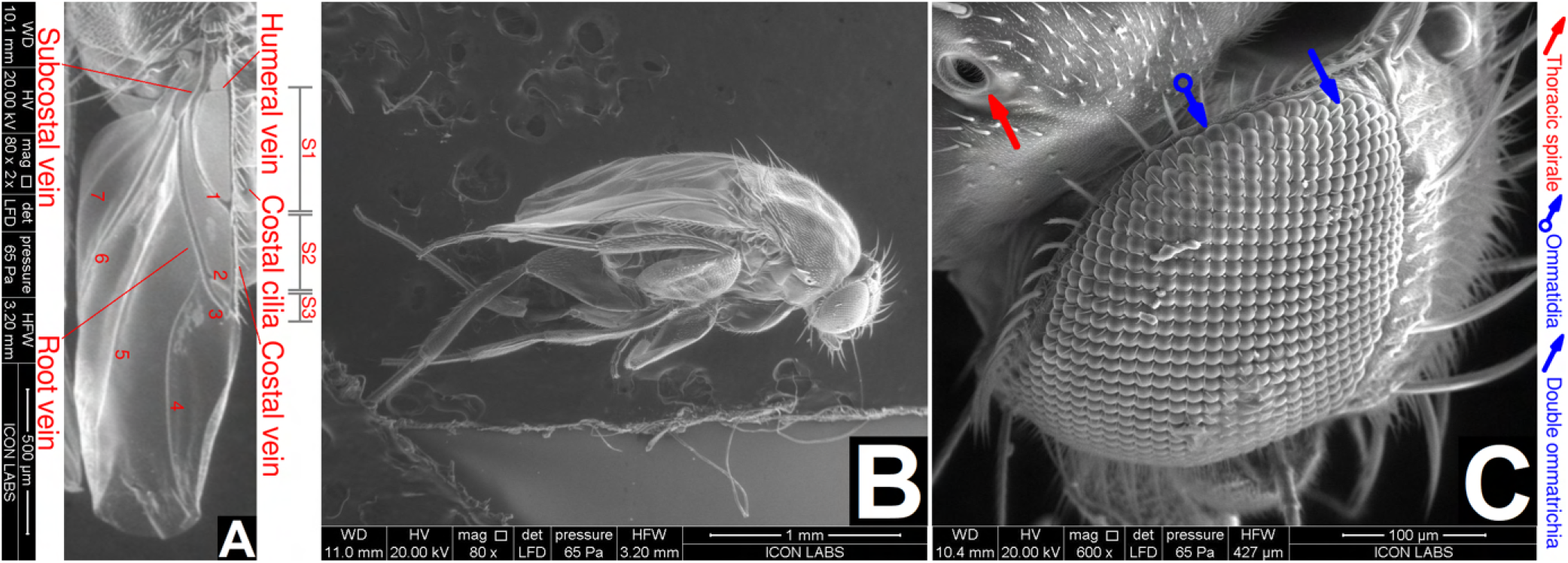
ESEM characterization of adult male *M. scalaris*: wings and lateral view. **(A)** A wing at 80× magnification displaying venation characteristic of scuttle flies. This panel has been digitally magnified by 2× for clarity. The costal, subcostal, humeral, and root vein are highlighted. Veins 1-7 and subcostal regions S1-S3 are also highlighted. Costal cilia can also be seen. textbf(B) Lateral view at 80× magnification. **(C)** Lateral view of the anterior cephalon at 600× magnification. The eye is the most prominent feature. Ommatidia (blue arrow) and double ommatrichia (blue arrow, hollow end) are highlighted. Note the square packing of the ommatidia. A thoracic spiracle (red arrow) is also observed

The other anatomical features of the head are typical of other phorid flies^2^. The head possesses numerous bristles. Ocellar bristles, postero-lateral bristles, pre-ocellar bristles, medio-lateral bristles, antero-lateral bristles, antial bristles, and supra-antennal bristles were all observed (Figure 11B). The antennae is 6-segmented. The segments (from proximal to distal) are as follows: scape, pedicel, flagellum, segment 4, segment 5, and the arista. The first two segments (scape, pedicel) are obscured. The flagellum is large and bulbous compared to that of *D. melanogaster*^34^. We hypothesize that this enlarged segment may contain sensitive sensory apparatus that enables *M. scalaris* to rapidly locate carrion, even if the carrion is concealed or difficult to access. The arista is elongated and branched. The ultrastructure of segments 4-5 have been described in more detail by Sukontason *et al*.^36^. The male mouthparts consist of a pair of vibrissae (Figure 11B), and are less developed than female mouthparts (Figure 14C). Female mouthparts contain vibrissae in addition to an oral opening. The labrum and labium are visible. Sukontason *et al*. have described the mouthparts in greater detail^37^.

From a dorsal view, the thorax appears smooth and densely covered with fine hairs (microchaete) (Figure 12B, 13A). The thorax also possesses large identifiable bristles (macrochaete) like those found on *D. melanogaster*^34^. The humeral, presutural, anterior supre-alar, posterior-supra alar, anterior post-alar, posterior post-alar, anterior scutellar, and posterior scutellar bristles were observed on the thorax (Figure 13C,D). The thorax possesses a pair of wings, three pairs of jointed legs, and two laterally placed thoracic spiracles.

**Figure 13.**
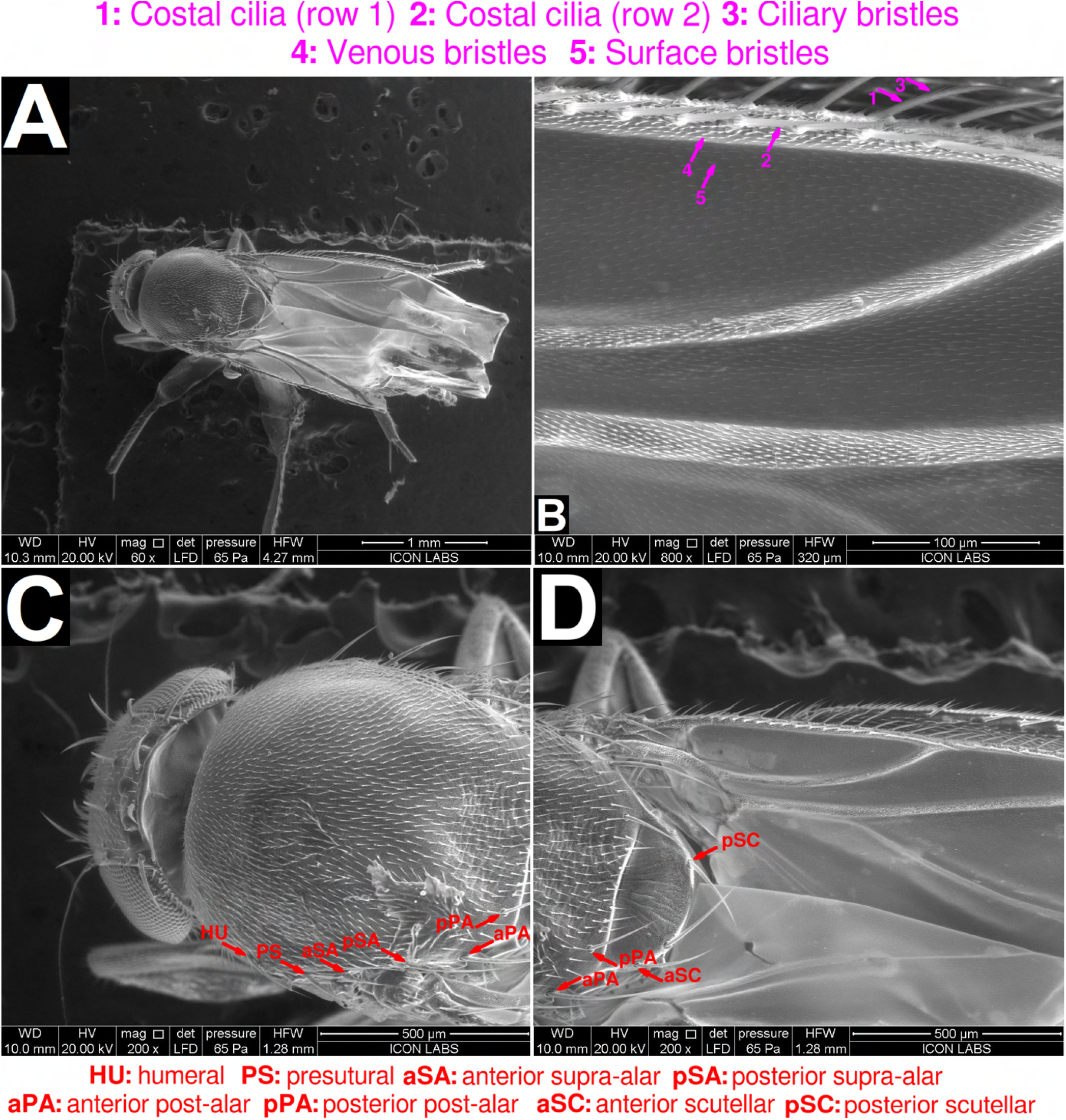
ESEM characterization of adult female *M. scalaris*. **(A)** Dorsal view at 60× magnification showing the head, thorax, and wings (not labelled). The distal portions of the wings were damaged while handling. **(B)** Dorsal view of the left wing at 800× magnification. Two rows of costal cilia (1,2) were observed. Ciliary bristles (3), venous bristles (4) and surface bristles (5) were also observed. **(C**,**D)** Dorsal view of the thorax, displaying humeral (HU), presutural (PS), anterior supra-alar (aSA), posterior supra-alar (pSA), anterior scutellar (aSC), and posterior scutellar (pSC) bristles (macrochaete). Smaller hairs (microchaete) can be seen covering the thorax (not labelled).

The wings possess venation (Figure 12A) characteristic of scuttle flies^2^. The costal vein extends out from the root along the anterior end of the wing. It is bordered by two rows of costal cilia (Figure 13B). The costal cilia themselves possess fine bristles, which we term as the ciliary bristles. Venous bristles (covering the veins) and surface bristles (covering all other surfaces) were observed. The subcostal vein, humeral vein, and root vein were also observed. Veins 1, 2, and 3 divide the wing into subcostal regions S1, S2, and S3, respectively. Veins 4, 5, 6, and 7 emerge from the root vein and extend to the wingtip. Axillary bristles were not observed.

The leg morphology of *M. scalaris* closely resembles that of *D. melanogaster*. Three pairs of legs are present on the thorax: prothoracic legs, mesothoracic legs, and metathoracic legs (Figure 11A). All legs possess the same segmentation pattern (from proximal to distal): coxa, trochanter, femur, tibia, and tarsus (Figure 11E). The tarsus is comprised of five tarsomeres. Two claws and two tentacled appendages are present on the distal end of each leg (Figure 11F, Figure 14D). We hypothesize that the claws are used to grip rough surfaces, while the tentacled appendages are used to grip smooth surfaces. *et. al* reported observing a sex comb on the male legs^38^. However we did not observe any such structure. We also observed long barbs projecting outward from the distal tibia.

**Figure 14.**
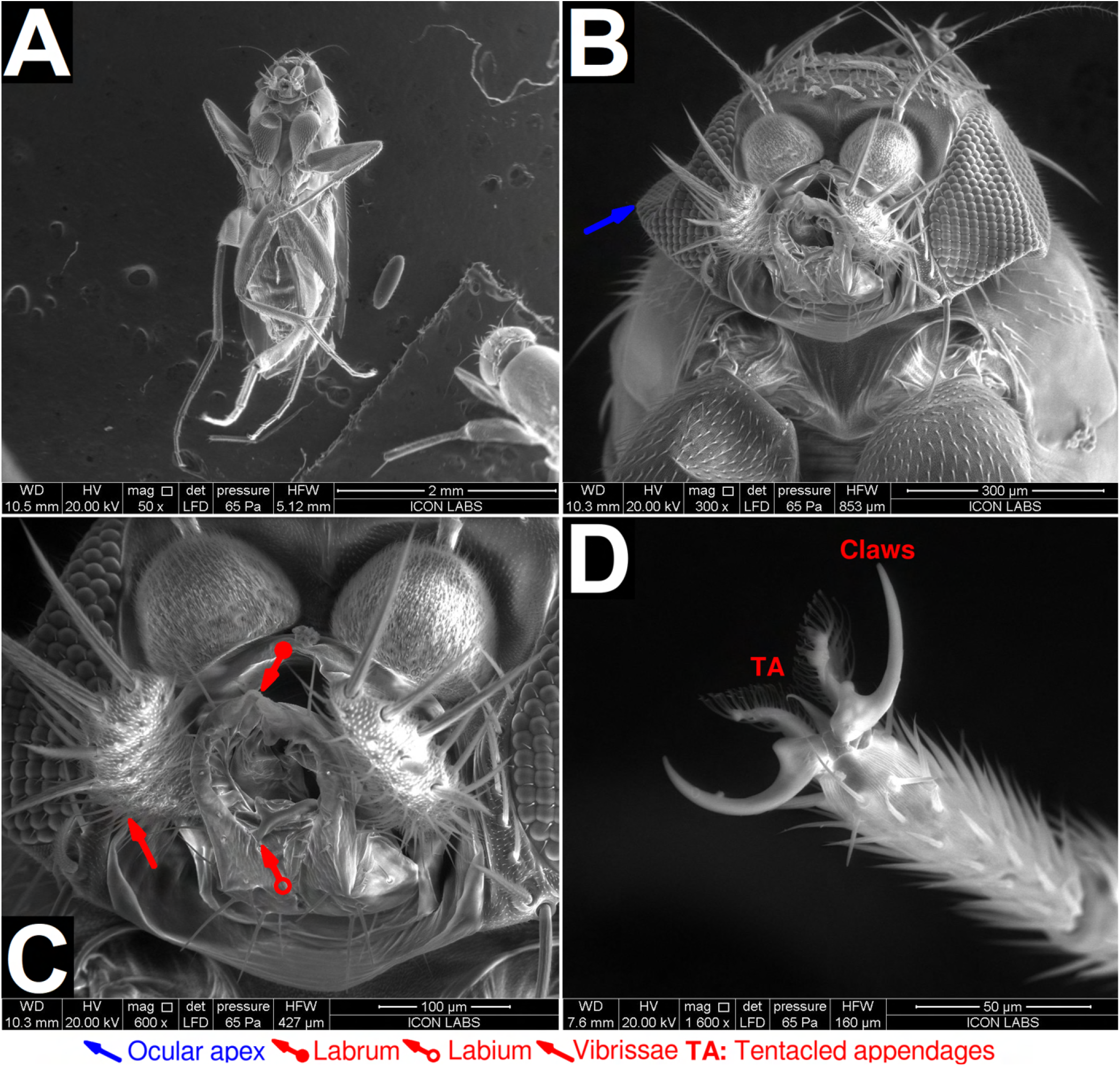
ESEM characterization of adult female *M. scalaris*. **(A)** Ventral view at 50× magnification showing the head, thorax, and abdomen (not labelled). **(B)** Wiew of the anterior cephalon at 300× magnification. Note the ocular apex (blue arrow). **(C)** View of mouthparts. The labrum (red arrow, round end), labium (red arrow, hollow end) and vibrissae (red arrow) are visible. **(D)** The distal end of a single metathoracic leg at 1600× magnification, showing the claws and tentacled appendages (TA).

*M. scalaris* bears characteristic black stripes on every segment of the dorsal abdomen (Figure 10A,D). Sex organs are present on the posterior end of the abdomen. The male possesses an intensely pigmented penis (Figure 10B,C,J), while the female possesses a relatively less pigmented ovipositor (Figure 10E,F). Zuha *et. al* have described the genitalia in greater detail^39^. A pregnant female is pictured in Figure 10G. Note the distended abdomen. A female immediately post-ovipositon is pictured in Figure 10H. Note the distended ovipositor and reduction in abdominal size.

During the process of image collection via light microscopy, we noticed an acephalic, or ‘headless’, phenotype in males (Figure 10I, J). 4 acephalic flies were observed in a cohort of 50 photographed males, indicating that the frequency of occurrence of this phenotype is 8%. We speculate that head may be damaged during the process of carapogenesis, during which the precarapace surrounding the pupal cephalon is ecdysed before the formation of the carapace. The pupal head is exposed and vulnerable to mechanical damage during this process. Alternatively, as all the flies possessing this phenotype were male, the trait may be linked to autosomal male determinants. Further study is warranted to explore the genetic and developmental processes that lead to this phenotype.

## Discussion

*M. scalaris* is a phorid fly that is an emerging model organism in the fields of developmental biology and genetics^8^. It is relevant in the field of forensic entomology for determining post-mortem intervals (PMIs) of corpses^15,16^. It is also clinically relevant as an etiological agent causing myasis^17–21^. Despite its relevance across fields, the morphological and ultrastructural characterization of *M. scalaris* remained fragmentary and incomplete, with reports from different workers characterizing no more than one life stage under different conditions.

In this work we report the first complete morphological characterization of *M. scalaris*. Using light microscopy (LM) and environmental scanning electron microscopy (ESEM), we have characterized the egg, all larval instars, pupal, and adult life stages of the fly. The comprehensive documentation of the morphological characteristics of all life stages of *M. scalaris* should serve as an ‘atlas’ to the community, allowing for the easy identification and comparison of different *M. scalaris* specimens.

We have described previously unreported morphological features and developmental processes. We described the sublabial flabellum, a ‘curtain’ that helps the larval oral cavity to open and close. We described the presence of oral ridge patterns that are unique to every larvae. We described pupal carapace and the process of carapogenesis: a process that involves the ecdysis of the dorso-anterior pupal crysalis (the precarapace) and its subsequent regrowth. We described the presence of an ocular apex,a ridge breaking the curvature of the compound eyes, on both eyes of male and female flies. Finally, we also reported the presence of an acephalic or ‘headless’ phenotype in adult male flies. Further work studying this phenotype could yield insights into the process of cephalogenesis.

## Methods

### Isolating *M. scalaris*

*M. scalaris* specimens were isolated from Malabar Hill, Mumbai, India (pin code: 400006). Flies were isolated using Whiskas catfood (adult, chicken in gravy). A pouch of catfood was placed in an exposed bowl in the area mentioned. Larvae were collected and used for subsequent experiments. This culture is currently maintained in our laboratory. Workers interested in acquiring our culture can contact the corresponding author.

### Culturing *M. scalaris*

Flies were subcultured using catfood media of the following composition: 75% catfood (homogenized using a mortar and pestle), 24% distilled water, 1% agar. Vials were incubated at room temperature. This media was autoclaved and 10 mL aliquots were poured into standard fruitfly vials. Culturing *M. scalaris* requires meat, and catfood was the most convenient source because: (1) catfood is sold in small packs of 85g, making it convenient for preparing small batches of vials; (2) unopened catfood can be stored at room temperature without the need for refrigeration; (3) catfood is processed into small, chewable cubes, minimizing the need for extensive homogenization; and (4) the composition of catfood from the same manufacturer will remain constant over different batches.

### Light microscopy

A trinocular stereo microscope with an attached camera (SHSM-1007 TDLED, Ski-Hi Optical Instruments) was used for this study. All specimens were placed on a hemocytometer for scale. All images were acquired at 20-40× magnification.

### Scanning electron microscopy

Environmental scanning electron micoscopy (ESEM) was performed using the FEI Quanta 200 Scanning electron microscope at Icon Labs Pvt. Ltd., Mumbai. Deceased larvae were used for ESEM after treatment with 70% isopropanol. Deceased adults were used for ESEM after exposure to a lethal dose of chloroform. Live eggs were used for ESEM. No fixation was necessary prior to performing ESEM.

## Acknowledgements

We would like to thank Kiran Rambhau Bhotkar (Assistant Manager - Application Support, Icon Labs Pvt. Ltd., Mumbai) for his excellent work as our SEM technician. His dedication, expertise, and attention to detail were instrumental in collecting electron micrographs for our samples.

## Author contributions statement

Must include all authors, identified by initials, for example: D.N. conceived the experiments, J.P, H.S, D.S, M.H, L.C, K.J, R.K, S.V. T.S., N.M, A.M, A.J, N.D., and P.S conducted the experiments, D.N. analysed the results. All authors reviewed the manuscript.

## Supplementary material

**Table S1** 16s rRNA sequences of *M. scalaris* larvae isolated in Mumbai, India.

**Table S2** Cytochrome c oxidase subunit I (COI) sequences of *M. scalaris* larvae isolated in Mumbai, India.

