## Supplementary Tables S1,S2 for "A complete morphological characterization of all life stages of the phorid fly *M. scalaris*"

### Supplementary material

| Gene | Primer name | Direction | Primer sequence |
|---|---|---|---|
| 16S rRNA | 16sSbr | Forward | CCGGTCTGAACTCAGATCACGT |
| **PCR product**<br>GGACTTTAAGTTTAAGCTGCTGCACCTAAAACTGTATCTTAATCCAACATCGAGGTCGCAATCTTTTTTATCAATATGAA<br>CTCTCTAAAAAAATTACGCTGTTATCCCTAAAGTAACTTGATTTCTTAATCATTAATAATGGATCAATAATTCATTAATT<br>TATGTTTATTATAAATTAAAAGTTTAACAAATTTTAATATCACCCCAATAAAATATATAAATTTATAATAATTTAATTTA<br>TCTATATAATTAAAATAATTTATATATAAAGATTTATAGGGTCTTCTCGTCTTTTAAAATTATTTTAGCTTTTTAACTAA<br>AAAATAAAATTCTATTATAAATTTATATGAAACAGTTAATATTTCATCCAACCATTCATACCAGCCTTCAATTAAAAGAC<br>TAATGATTATGCTACCTTTGCACAGTTAAGATACTGCGGCCATTTAAAAAAATTTCAGTGGGCAGGCTAGACTTTAAATTA<br>AATTCAAAAAGACATGTTTTTGATAAACAGGCGA | | | |
| **Top-scoring hit (BLAST)** | **Species** | **Percent identity** | **E value** |
| NC_023794.1 | *Megaselia scalaris* | 99.61% | 0.0 |

| Gene | Primer name | Direction | Primer sequence |
|---|---|---|---|
| 16S rRNA | 16sSar | Reverse | CGCCTGTTTATCAAAAACAT |
| **PCR product**<br>TATATTTAAAGTCGAGCCTGCCCACTGAGATTTTTTAAATGGCCGCAGTATCTTAACTGTGCAAAGGTAGCATAATCATT<br>AGTCTTTTAATTGAAGGCTGGTATGAATGGTTGGATGAAATATTAACTGTTTCATATAAATTTATAATAGAATTTTATTT<br>TTTAGTTAAAAAGCTAAAATAATTTTAAAAGACGAGAAGACCCTATAAATCTTTATATATAAATTATTTTAATTATATAG<br>ATAAATTAAATTATTATAAATTTATATATTTTATTGGGGTGATATTAAAATTTGTTAAACTTTTAATTTATAATAAACAT<br>AAATTAATGAATTATTGATCCATTATTAATGATTAAGAAATCAAGTTACTTTAGGGATAACAGCGTAATTTTTTTAGAGA<br>GTTCATATTGATAAAAAAGATTGCGACCTCGATGTTGGATTAAGATACAGTTTTAGGTGCAGCAGCTTAAACTTAAAGTC<br>TGTTCGACTTTTAAATTCTTACGTGATCTGATTTCAGACCGGATG | | | |
| **Top-scoring hit (BLAST)** | **Species** | **Percent identity** | **E value** |
| NC_023794.1 | *Megaselia scalaris* | 99.04% | 0.0 |

**Table S1.** 16S rRNA PCR products of *M. scalaris* larvae isolated in in Mumbai, India.

| Gene | Primer name | Direction | Primer sequence |
|---|---|---|---|
| COI | COI_LCO1490 | Forward | GGTCAACAAATCATAAAGATATTGG |
| **PCR product** | | | |
| TTGATTTATTTTTGGAGCCTGAGCTGGAATAGTAGGAACATCTTTAAGTATTATAATTCGAGCTGAATTAGGGCACCCCG | | | |
| GTGCTTTAATTGGTGATGATCAAATTTATAATGTAATTGTTACTGCCCATGCATTTATTATAATTTTTTTTATAGTAATA | | | |
| CCTATTATAATAGGAGGATTTGGAAATTGATTAGTTCCCTTAATATTAGGGGCACCTGATATGGCTTTTCCACGAATAAA | | | |
| TAATATAAGTTTTTGAATACTTCCCCCTTCTCTAACTCTTTTATTAGCAAGAAGTATAGTAGAAAATGGAGCCGGAACTG | | | |
| GTTGAACAGTTTATCCACCCCTATCTTCTAGAATTGCCCATAGAGGAGCTTCAGTCGATTTAGCAATTTTTTCATTACAT | | | |
| CTTGCCGGAATTTCTTCTATTCTTGGAGCAGTAAATTTTATTACTACAATTATTAATATACGATCTACAGGAATTACTTT | | | |
| TGATCGAATACCTTTATTTGTATGATCAGTAGGTATTACTGCTCTTTTATTATTACTTTCACTACCTGTTCTAGCAGGTG | | | |
| CTATTACTATACTATTAACAGACCGAAATTTTAATACATCATTCTTTGATCCTGCGGGAGGGGGAGATCCAATTCTATAT | | | |
| CAACATTTATTTTGATTTTGGGGGCCATCCAAA | | | |
| **Top-scoring hit (BLAST)** | **Species** | **Percent identity** | **E value** |
| KX832638.1 | *Megaselia scalaris* | 99.54% | 0.0 |

| Gene | Primer name | Direction | Primer sequence |
|---|---|---|---|
| COI | COI_HCO2198 | Reverse | TAAACTTCAGGGTGACCAAAAAATCA |
| **PCR product** | | | |
| GTGCATGGATTGGATCTCCCCTCCTGCAGGATCAAAGAATGATGTATTAAAATTTCGGTCTGTTAATAGTATAGTAATAG | | | |
| CACCTGCTAGAACAGGTAGTGAAAGTAATAATAAAAGAGCAGTAATACCTACTGATCATACAAATAAAGGTATTCGATCA | | | |
| AAAGTAATTCCTGTAGATCGTATATTAATAATTGTAGTAATAAAATTTACTGCTCCAAGAATAGAAGAAATTCCGGCAAG | | | |
| ATGTAATGAAAAAATTGCTAAATCGACTGAAGCTCCTCTATGGGCAATTCTAGAAGATAGGGGTGGATAAACTGTTCAAC | | | |
| CAGTTCCGGCTCCATTTTCTACTATACTTCTTGCTAATAAAAGAGTTAGAGAAGGGGGAAGTATTCAAAAACTTATATTA | | | |
| TTTATTCGTGGAAAAGCCATATCAGGTGCTCCTAATATTAAGGGAACTAATCAATTTCCAAATCCTCCTATTATAATAGG | | | |
| TATTACTATAAAAAAAATTATAATAAATGCATGGGCAGTAACAATTACATTATAAATTTGATCATCACCAATTAAAGCAC | | | |
| CGGGGTGCCCTAATTCAGCTCGAATTATAATACTTAAAGATGTTCCTACTATTCCAGCTCAGGCTCCAAAAATAAAATAT | | | |
| AAAGTTCCAATATCTTTATGATTTGGTTGACCAAA | | | |
| **Top-scoring hit (BLAST)** | **Species** | **Percent identity** | **E value** |
| MT396301.1 | *Megaselia scalaris* | 99.40% | 0.0 |

**Table S2.** Cytochrome c oxidase subunit I (COI) PCR products of *M. scalaris* larvae isolated in in Mumbai, India.
